# Antibiotic-induced Recombination in *Escherichia coli* Requires the Formation of DNA Double-Strand Breaks

**DOI:** 10.1101/2022.03.08.483535

**Authors:** Arpita Nath, Dan Roizman, Israa Amin, Nivetha Pachaimuthu, Jesús Blázquez, Olga Makarova, Jens Rolff, Alexandro Rodríguez-Rojas

## Abstract

Recombination is an essential process in bacterial drug resistance evolution. Fluoroquinolones, an important class of antibiotics, are known to stimulate recombination in *Escherichia coli*. When bacteria are exposed to antibiotics other than fluoroquinolones, including strong up-regulators of recombination pathways, there is no detectable change in the recombination level. Here we explore why fluoroquinolones, but not other antimicrobials, increase recombination rates. Fluoroquinolones, in contrast to other antibiotics, generate DNA double-strand breaks (DSBs). We tested whether other drugs that also cause double-strand breaks, such as mitomycin C and bleomycin, also affect bacterial recombination rates with consistent increases in recombination. A positive correlation between the number of DSBs and the recombination frequency was found. The manipulation of the level of DSBs directly impacted the recombination frequency. Our results highlight that only antibiotics that induce DNA double-strand breaks are more probable to increase genetic diversity via recombination. The stimulation of recombination by DSB-causing antimicrobials is an additional factor leading to the risk of antibiotic resistance evolution.

## Introduction

A recent study provides the most comprehensive global assessment to date of bacterial antibiotic resistance (AMR). It covers 204 countries and over 520 million data records. In 2021, 4.71 million deaths were linked to AMR, 1.14 million of which were direct. Projections indicate that without increased control measures, AMR could cause up to.9 million direct deaths annually by 2050 (1). The number of deaths varied between regions, with the highest toll being reported in regions with lower-resource settings. *Escherichia coli, Staphylococcus aureus, Klebsiella pneumoniae, Streptococcus pneumoniae, Acinetobacter baumannii*, and *Pseudomonas aeruginosa* were the six most significant pathogens leading to antimicrobial resistance (2). Resistance to fluoroquinolones and β-lactams, the first-line antibiotics in most infections, contributed to at least 70% of AMR deaths (3). Antibiotics are a selective pressure in themselves, but they also enhance evolvability by elevating mutation rates and account for increased recombination frequency and lateral gene transfer in bacteria (4).

An excellent example of an increasing issue with antibiotic resistance over the years is the quinolones, to which resistance emerges as soon as they are introduced (5, 6). The mechanism of quinolone action is the inhibition of two essential bacterial type II topoisomerase enzymes, DNA gyrase and DNA topoisomerase IV (7). Both of these enzymes are hetero-tetramers with two subunits. The activity of both enzymes causes the catalysis of a DNA double-strand break (DSB), passing another DNA strand through the break and then resealing it together to regulate the DNA supercoiling state. DNA strand passing occurs via domains localized in GyrA and ParC, whereas the ATPase activity is required for enzyme catalysis and is located in domains belonging to GyrB and ParE subunits (8).

The SOS stress pathway is activated when several stressors induce DNA damage; this includes sub-lethal antibiotic treatments. Consequently, error-prone alternative DNA polymerases repair the damage at the price of introducing mutations, a process called translesion synthesis (9). The general stress response controlled by the sigma factor RpoS in Gram-negative bacteria also increases mutation rates under antibiotic stress by indirectly down-regulating MutS, a key protein of the DNA mismatch-repair system (10). Moreover, other classes of antibiotics, such as sulfonamides, increase mutagenesis directly by creating an imbalance in the nucleotide pool during replication (11). All these direct or indirect mutagenic effects are common to many families of antimicrobials except for cationic antimicrobial peptides that do not increase the mutation rate or recombination, probably because they do not induce mutagenic pathways such as the SOS or RpoS stress responses (12, 13).

Recombination is a fundamental driver of antimicrobial resistance by inducing, for example, gene duplication and amplification (14, 15). It is known that fluoroquinolones cause an increase in the recombination rate in *E. coli*, suggesting the potential for acquiring antibiotic resistance (16). In pathogenic *E. coli* isolates, recombination rates can be significantly higher than in commensal strains (17). However, the number of antibiotics that increase recombination in *E. coli* is so far limited to fluoroquinolones, suggesting this is a unique property of this family (18).

Here we asked why fluoroquinolones, but not other antimicrobials, increase bacterial recombination rate. Many antibiotics induce the SOS system, upregulating several recombination proteins such as RecA. Nevertheless, this overexpression alone does not increase recombination. Previous data suggests that the induction of DSBs, a hallmark of fluoroquinolone-induced damage, could be responsible for boosting the recombination rate (16). Therefore, we hypothesized that to stimulate recombination by antibiotics in Gram-negative bacteria such as *E. coli*, the induction of the SOS system alone is not sufficient and that generating DNA double-strand breaks is necessary. To assess the generality of our findings, we compared ciprofloxacin to other non-fluoroquinolone drugs known for their capacity to generate DSBs: mitomycin C (19) and bleomycin (20).

## Results and Discussion

First, we measured the recombination frequency of several antimicrobials using a plasmid-based system described elsewhere (Figure S1) (17). Since the plasmid is a multi-copy system, it has the advantage of being more sensitive (17) than other chromosomal integrated systems used before. We used the minimal inhibitory concentration (Table S1) as a reference value for the experiments with antimicrobials that covered diverse antibiotic classes. This experiment confirmed that from the initially tested panel, only ciprofloxacin increased recombination rates, which is in agreement with previous reports for ciprofloxacin (13, 16, 18). What makes ciprofloxacin different from other antibiotics is its ability to induce DSBs, which we thought was the main cause of the observed increase in recombination. We therefore also tested other antimicrobials that can induce this type of DNA damage such as mitomycin C (19) and bleomycin (20). In our experiments, we also included another antibiotic of diverse families. Interestingly, we found that mitomycin C and bleomycin significantly increased the recombination rate by more than one order of magnitude compared to the controls’ basal level and similar to ciprofloxacin (Figure 1). These results provide evidence that our hypothesis indicating that DSBs mediated the increase of recombination rate by fluoroquinolones is correct.

**Figure 1.**
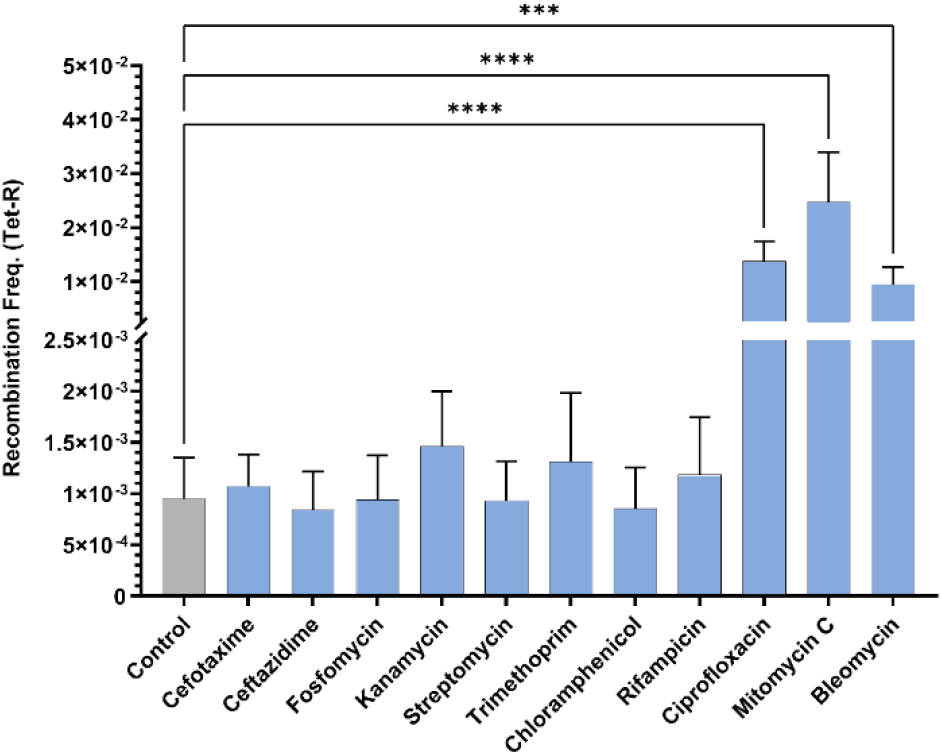
Recombination frequencies of *E. coli* MG1655 harboring the plasmid pRHOMO (recombination construct) treated with antibiotics at their 1xMIC for three hours. Bars represent the respective means of five biological replicates and their standard deviation (n=5 + SD) values of recombination frequency of the bacteria. Differences in recombination frequencies between treatments were analyzed by an ordinary one-way ANOVA followed by Dunnett’s multiple comparisons tests with a single pooled variance, performed using GraphPad Prism 9 (*p*-values denote: *** - <0.0005, **** - <0.0001). Ciprofloxacin, mitomycin C and bleomycin had a significantly higher recombination frequency compared to the control group.

The recombination frequency calculation in this study relies on the number of tetracycline-resistant colonies of *E. coli* harboring the pRHOMO plasmid that appears in the LB plates upon antibiotic treatments (see Material and Methods section for detailed description). Ciprofloxacin and other antimicrobials used in this work can increase mutagenesis (21, 22). One possible bias is the risk of spontaneous resistant mutants to tetracycline, which could be scored as recombination frequency increases. We treated *E. coli* MG1655 with the same concentration of antibiotics and experimental conditions, and we did not obtain any colony by plating the same inoculum and a ten-fold higher one. These results ensure that the colonies resistant to tetracycline were generated only due to recombinational events (the reconstitution of the tetracycline-resistant cassette) and not due to spontaneous mutations.

Mitomycin C and bleomycin also induce the SOS response as they cause direct DNA damage (20, 23). To understand the relevance of the SOS system in the observed higher recombination phenotype, we used a RecA mutant unable to trigger the SOS response and lacking the most important protein for DSB repair in *E. coli*. We found that for ciprofloxacin, mitomycin C and bleomycin, the increase in recombination frequency was dependent on the recombination protein RecA (Figure 2). A RecA-deficient mutant suppressed the recombination increase to levels comparable to the controls. This experiment, along with our previous data, suggests that the recombination stimulation requires RecA protein and DSBs, indicating that the first step is DNA breakage, and the second step is damage repair. The activation of SOS, which regulates RecA, is not always correlated with the presence of DSBs, but DSBs always seem to trigger the transcription of SOS genes in our experiments. In other words, the SOS system components can increase the recombination rate but its activation mayt not be necessary (24). Indeed, it has been reported in previous works that the increase of recombination rate by ciprofloxacin does not require SOS activation (16).

**Figure 2.**
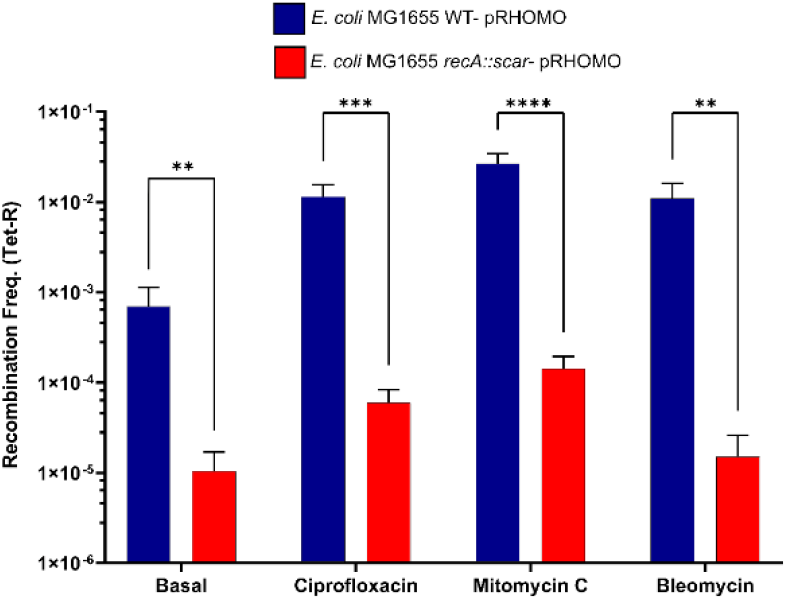
Effect of RecA inactivation on recombination frequency induced by antibiotics that promote double-strand breaks. Antibiotic-stimulated recombination for mitomycin C, bleomycin, and ciprofloxacin (1xMIC during three hours) is primarily dependent on RecA protein repair activity since *E. coli* MG1655 *recA*::scar harboring the plasmid pRHOMO suppressed the change in recombination rate observed in the wild-type strain when treated with DSB-inducer antimicrobials. Differences in recombination rates between strains were analyzed by unpaired two-tailed student t-tests, considering similar SD and 95% Confidence Intervals. Asterisks represent statistical differences (Basal: ** - *p*=0.0076, Ciprofloxacin: *** - *p*=0.0002, Mitomycin C: **** - *p*<0.0001, Bleomycin: ** - *p*=0.0015, respectively). Each experiment consisted of five replicates, including non-treated controls (basal recombination rate value of the non-treated bacteria). The bars represent the means, while the error bars represent the standard deviations (n=5 + SD).

To confirm that DSBs are significant determinants of the induction of recombination, we correlated the DSBs lesions and the recombination rate. For that purpose, we determined the number of DSBs for ciprofloxacin, mitomycin C and bleomycin. We used a reporter system (kindly gifted by Susan M. Rosenberg from Baylor College of Medicine, Texas, USA) that allows observing these lesions in real-time. The system consists of a green fluorescent protein (GFP) fusion containing bacteriophage GAM protein. GAM protein recognizes the DNA ends of DSBs, while GFP allows the tagging and forms foci (25). We found a positive correlation between the number of DSB lesions and recombination frequency (Figure 3). In complementary experiments, we repeated this analysis to check if there was a positive correlation between antibiotic concentration and recombination frequency. For this, we used ciprofloxacin at three different concentrations and determined the number of DSBs at each dose. We found that as the ciprofloxacin concentration increased, the recombination frequency also increased (Figure 4). Microscopic pictures of cells treated with growing concentrations of DSB-inducing antibiotics show the induction of DSBs by ciprofloxacin, mitomycin C and bleomycin (Figure 5). None of the other antibiotics used in this study produced DSBs, as indicated by the absence of GFP-Gam foci. This is another result indicating the fundamental role of DSBs in the ability of antibiotics to stimulate recombination.

**Figure 3.**
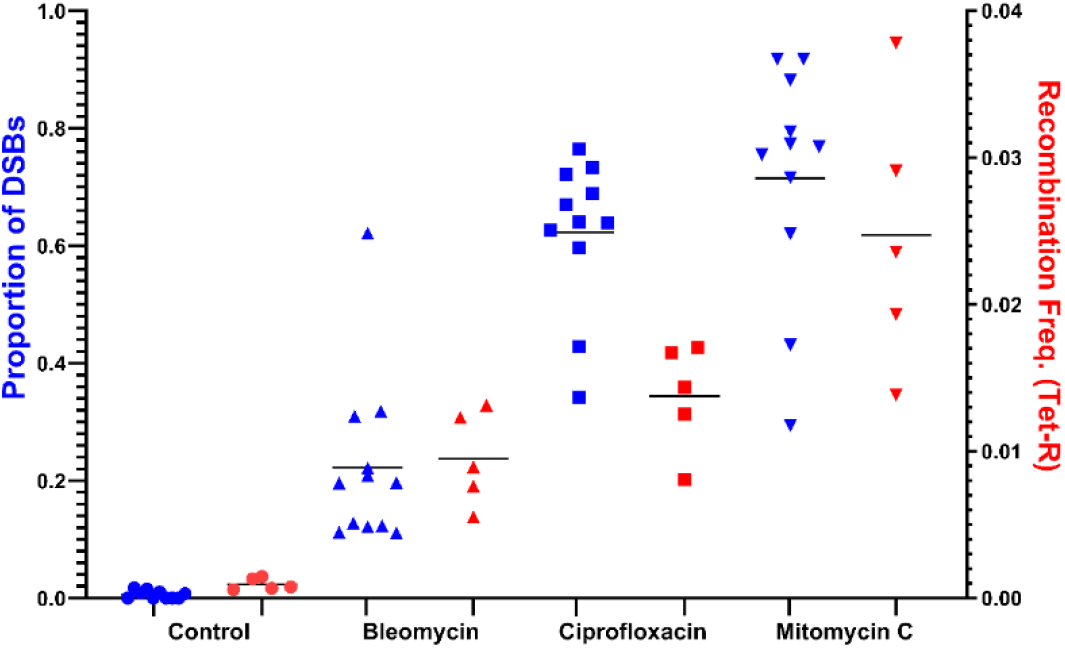
Co-occurrence of DSB and the increase of recombination rate for bleomycin, ciprofloxacin and mitomycin C. The graph shows a significant positive linear trend (Spearman correlation coefficient r= 0.819, *p*<0.0001, Slope=0.253 +/-0.0193) between the number of DSBs lesions (in blue) and increased recombination frequency (in red) when bacteria are treated with antibiotics that induce DSBs. Each experiment consisted of five replicates and had its non-treated control (basal recombination rate value of the non-treated bacteria, n=5). The positive control for DSBs (I-*SceI*) consisted of a unique restriction site in the bacterial chromosome that can be cut by inducing the expression of the restriction enzyme *SceI*.

**Figure 4.**
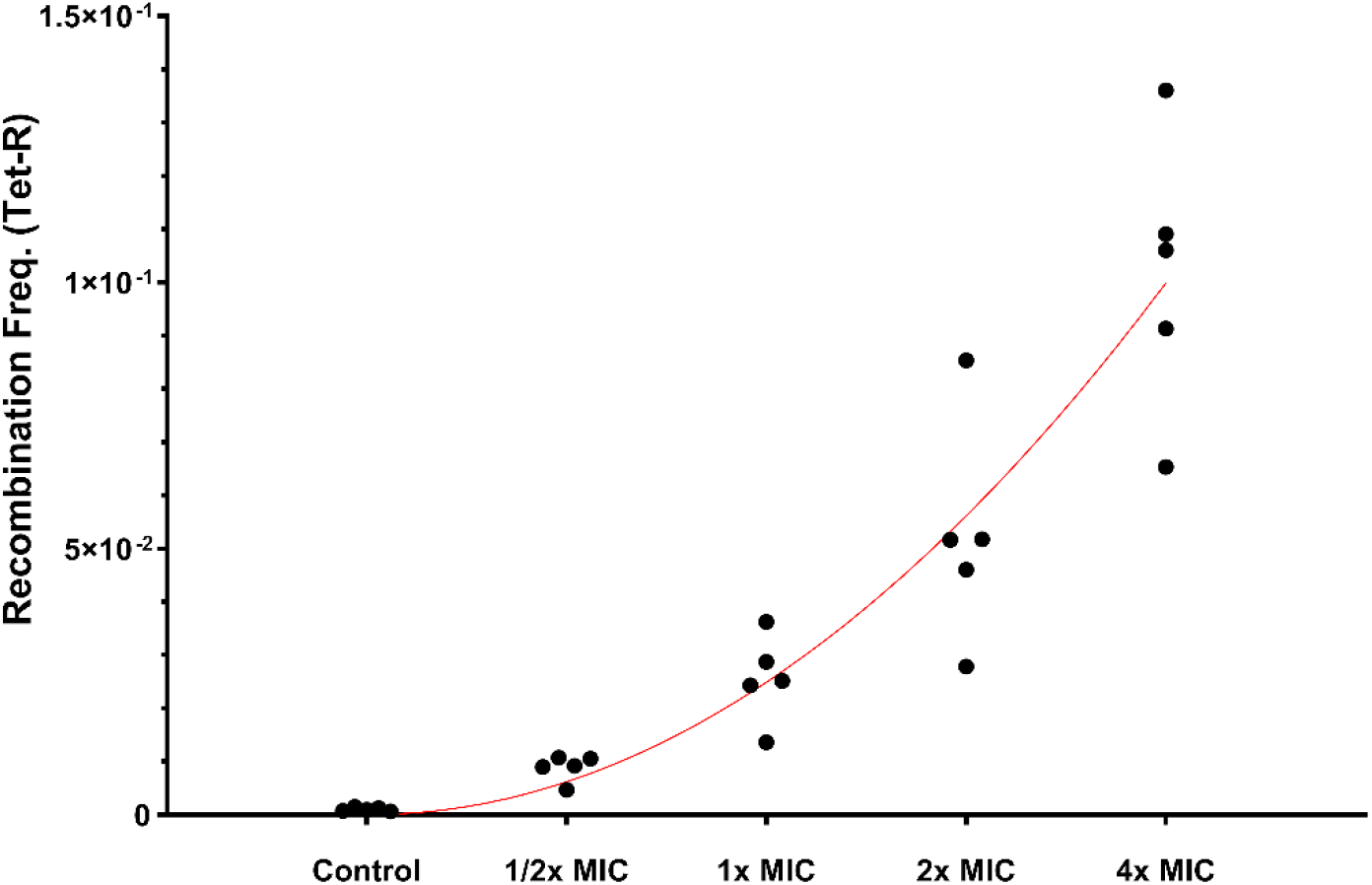
Dose-response of recombination frequency of *E. coli* MG1655-pRHOMO treated with growing ciprofloxacin. The graph shows a positive trend in the recombination frequency and increasing ciprofloxacin dose (Fit based on simple quadratic equation, Spearman correlation coefficient r=0.873, p-value<0.0001). Each experiment consisted of five replicates and had its non-treated control (basal recombination rate value of the non-treated bacteria, n=5).

**Figure 5.**
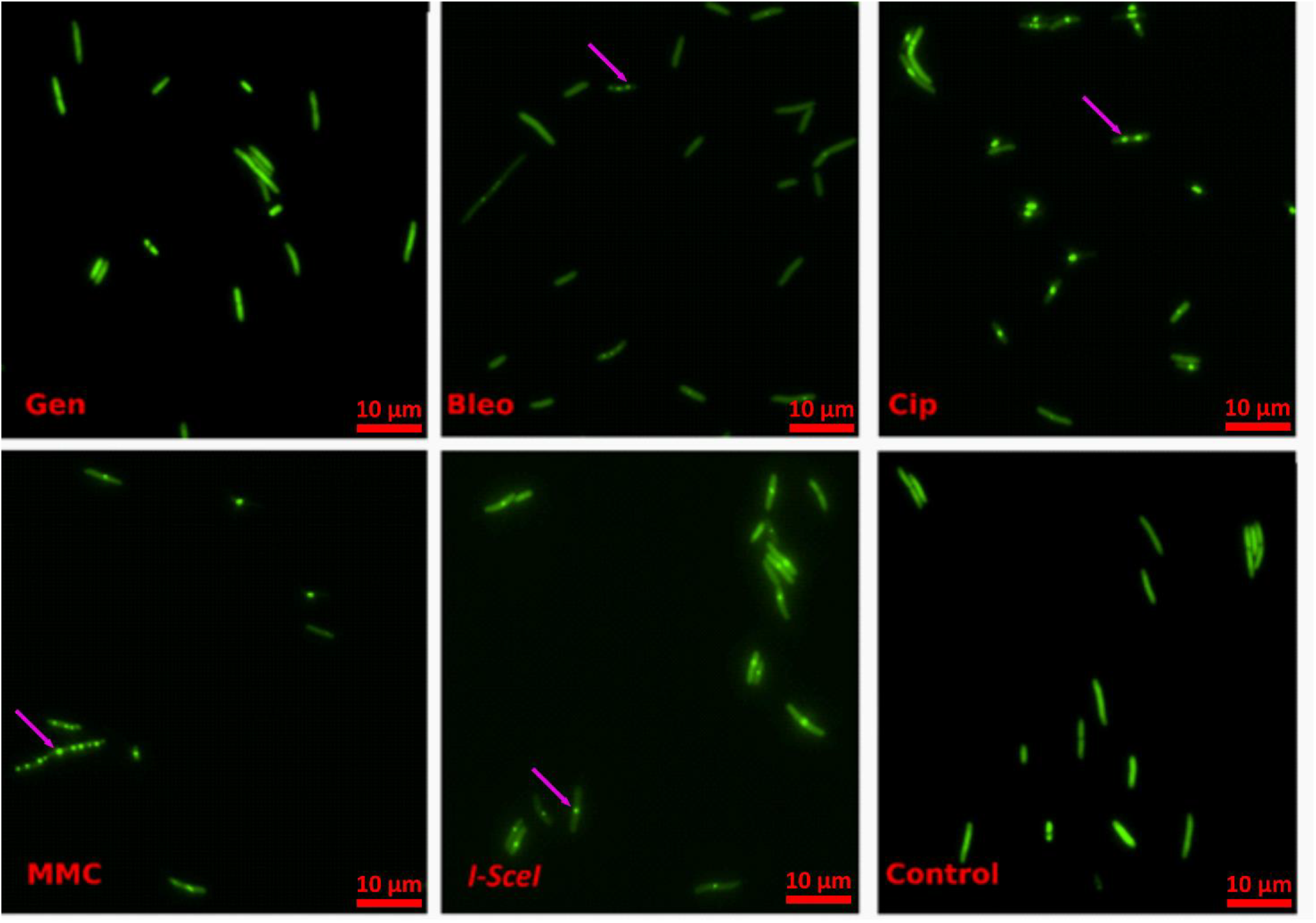
Fluorescent microscope images of antibiotic-generated double-strand breaks after treating the *E. coli* strain SMR14354 with gentamicin (Gen), bleomycin (Bleo) and ciprofloxacin (CIP), MIC of mitomycin C (MMC) with the minimal inhibitory concentration for 3 h. The green foci (indicated by magenta arrows) indicate the binding of the GFP-GAM fusion protein to free DNA ends and are only found at high frequency in treated bacteria with bleomycin, ciprofloxacin and mitomycin C. The foci formation (indicating the presence of DSBs) can also be observed in the positive control (I-*SceI*), which consists of a unique restriction site in the bacterial chromosome that can be cut by inducing the expression of the restriction enzyme *SceI*. Gentamicin is used as a control antibiotic, which does not increase recombination frequency or the number of DSBs.

Afterwards, we tested whether DSBs, caused by antibiotics, are responsible for antibiotic-induced recombination. One of the antibiotics that produce DSBs, bleomycin, requires iron as a cofactor to induce DSBs (20, 26). In the experiments described so far, we used LB, an iron-rich medium (27). Therefore, we reduced the iron content of the medium by suppressing iron (by adding 2–2′ bipyridyl) during the bleomycin treatment. Suppressing the iron content significantly decreased the recombination frequency (Figure 6), consistent with the hypothesis that DSBs cause changes in recombination. Complementarily, it is known that under low oxygen tension, the actual damage induced by bleomycin consists of forming oxidized abasic sites, while in aerobic environments, single and double-strand breaks predominate (28). The reason for the decrease in DSBs under anaerobic conditions is due to a decrease in Fenton reactions that do not require only iron but also oxygen-stimulated hydrogen peroxide production during respiration (29).

**Figure 6.**
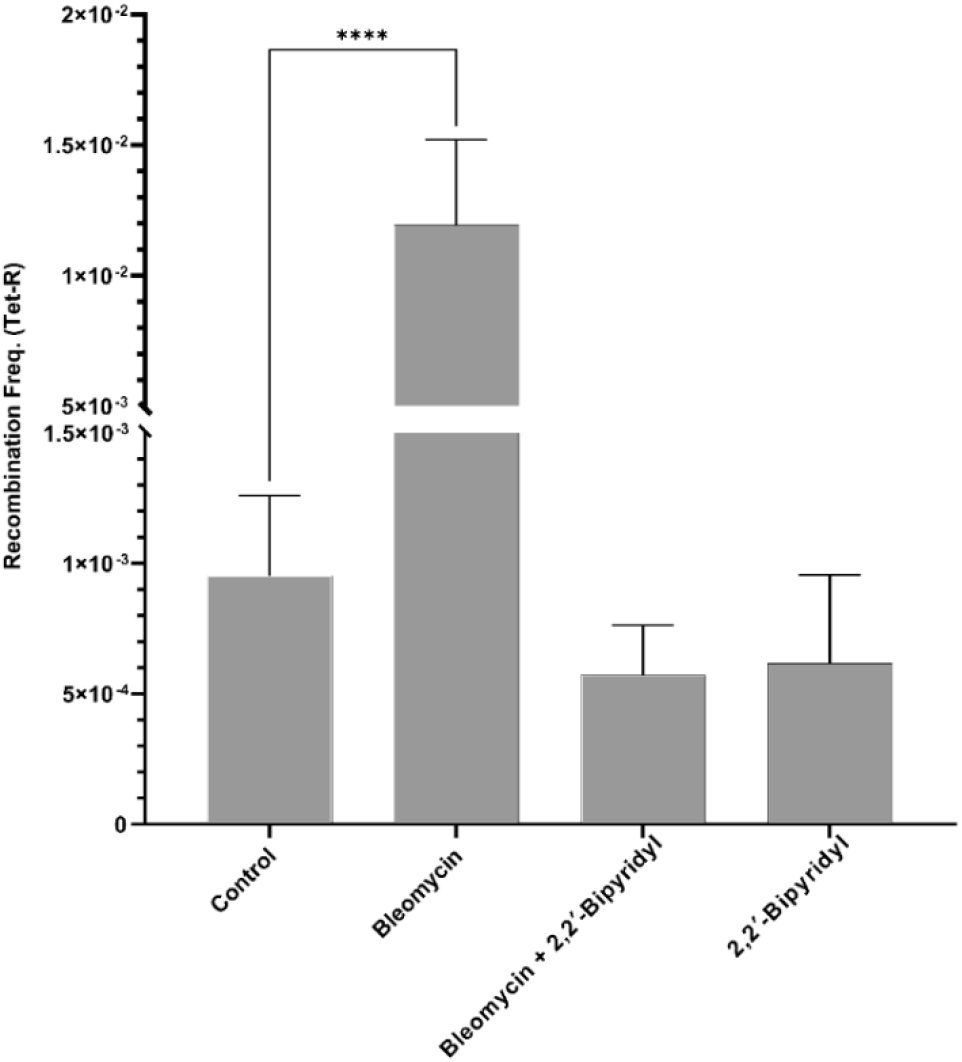
Impact of iron chelation by 2–2′ bipyridyl on recombination frequency of *E. coli* MG1655-pRHOMO treated with 1xMIC of bleomycin. LB medium contains between 15 to 20 µM of iron (27), which potentiates the capacity of bleomycin to induce DSBs (20) (Analyzed by an ordinary one-way ANOVA followed by Dunnett’s multiple comparisons tests with a single pooled variance, performed using GraphPad Prism 9, p-value denotes: **** - <0.0001). The experiment consisted of five independent replicates (n=5), bars representing the mean and standard deviation. A significant decrease in recombination rate was observed when an iron-titrating level of 2–2′ bipyridyl (final concentration of 50µM) was added together with bleomycin, leading to its return to basal level.

Our results point to an essential trait of recombination, at least in Gram-negative bacteria. The canonical recombination machinery of *E. coli* in the absence of stress is extremely conservative. Even in the presence of substrate for recombination, the frequency of recombination events is low and seems to be more important for repair processes. Consistent with this notion is the need to use helper recombination systems (e.g., λ red recombinase) to increase the probability of DNA integration into the chromosome through homologous recombination (30, 31). An additional implication of our result is that the induction of double-strand breaks and hence recombination by antibiotics could increase genetic diversity and enhance the risk for antimicrobial resistance. The phenotype of enhanced recombination rate could happen in natural scenarios since two different soil bacteria produce bleomycin and mitomycin C (32, 33). Some other antibiotics, such as novobiocin and nitrofurantoin, damage DNA and are produced by soil microorganisms (34), suggesting that the role of DSBs in promoting recombination as described here could be a ubiquitous phenomenon in environmental scenarios. Our study also reinforces the idea of controlling the pollution with other DNA damaging antibiotics such as fluoroquinolones that are incredibly stable and have been shown to accumulate in the environment (35).

We note that some antibiotics could produce a reduction in population or differential partial killing, and thus, an uneven number of cell divisions could impact the recombination frequency while comparing the treatments among themselves. For this reason, we did another experiment where we simulated this situation. We inoculated cultures of *E. coli* MG1655-pRHOMO, with inoculum sizes ranging from 10^4^ to 10^8^ bacteria (Figure S2). This experiment estimates the contribution of additional number of cell division to the recombination frequency. We did not find any detectable difference in recombination frequency due to additional number of divisions. This experiment simulates the uneven killing effect of different antibiotics since the starting number of bacteria could differ among treatments during recovery and after removing the antimicrobial treatments.

Taken together, our results provide compelling evidence that antibiotics that induce DSBs can increase the basal level of recombination in *E. coli*. This conclusion answers our central question on how ciprofloxacin could increase the recombination rate, at least in *E. coli*. Ciprofloxacin is a broad-spectrum antibiotic with a conserved action mechanism across diverse bacterial species (8, 36). Because the three antibiotics in the focus of this work are broad-spectrum and produce similar damage in other bacterial species, we can foresee these results being potentially extendable to other microbes.

The implications of our work can be extended to cancer research. Since bleomycin and mitomycin C are frequently used as antitumor drugs (28, 37), there is a chance that they would also increase gene shuffling in tumoral cells through recombination. Future studies could then uncover whether an increase in recombination rate is also possible in tumor cells, leading to chemotherapy resistance and carcinoma recidivation (38).

To further investigate whether oxidative stress contributes to the observed increase in recombination, we added thiourea and N-acetyl-L-cysteine, both reactive oxygen species (ROS) scavengers, to our assays. The presence of either thiourea or N-acetyl-L-cysteine did not significantly alter the recombination rate in ciprofloxacin-treated cultures at different doses (Supplementary Figure SX). This suggests that the stimulation of homologous recombination is not a consequence of oxidative stress. This finding supports the hypothesis that the crucial trigger is the induction of double-strand breaks (DSBs), rather than ROS accumulation or other indirect stress responses.

We also investigated the possibility that ciprofloxacin-resistant strains, which we had developed using the same genetic background (*E. coli* MG1655), would be less prone to recombination when treated with different antibiotic doses. In line with this view, we tested three ciprofloxacin-resistant derivatives which, as expected from the known mechanism of fluoroquinolone resistance, exhibited different mutations in target genes (gyrA, marA). No evidence of recombination-mediated gene exchange or rearrangement was found in these resistant isolates. In summary, these results confirm that ciprofloxacin increases the recombination frequency through its ability to induce DNA double-strand breaks (DSBs), not through enhanced mutagenesis or ROS-dependent stress. In contrast, treatments that do not induce DSBs, including those in combination with thiourea, do not lead to an increase in the recombination rate, reinforcing the causal relationship between DNA breaks and the activation of homologous recombination. Our results confirm that the stimulation of homologous recombination by ciprofloxacin is directly related to its ability to generate DNA double-strand breaks (DSBs) and not to general stress responses or oxidative damage. The addition of thiourea, a radical scavenger, had no effect on the recombination rate. The ciprofloxacin-induced increase in the recombination rate is not reduced, indicating that reactive oxygen species are not required for this effect. This observation supports the conclusion that the presence of DNA double-strand breaks (DSBs) is the main signal that triggers recombination under these conditions.

In accordance with previous results by López et al. (Mol. Microbiol., 2007), we interpret DSB processing via the recBCD and recFOR signaling pathways as essential for the observed increase in homologous recombination, whereas activation of the SOS response alone is insufficient to explain this effect. In line with this mechanism model, the ciprofloxacin-resistant lines obtained in our experiments exhibited specific point mutations in gyrA and parC that are characteristic of quinolone resistance, and no evidence of recombination-mediated genomic rearrangements. Therefore, the increased recombination frequency induced by DSB-inducing antibiotics such as ciprofloxacin reflects enhanced repair activity and not a direct pathway to resistance development.

Our results also have implications for bacterial evolution. It has been shown that recombination facilitates adaptive evolution, particularly in fluctuating environments that require complex genomic changes (39). For example, work in *E. coli* has shown that, in the presence of doxycycline and erythromycin, costly adaptive evolution is facilitated by recombination (38). Recent work in *pneumococci* demonstrated the importance of interspecies recombination in acquiring resistance against two antibiotics, tetracycline and macrolides (40).

Recombination in bacteria occurs by two principal mechanisms. By one side, an allelic exchange can buffer genetic variability within bacterial populations as gene exchange mediates selective sweeps. On the other hand, incorporating new genes via horizontal gene transfer or gene duplication through recombination leads to an increase in the size of the genome (41). Horizontal gene transfer, which frequently entails recombination events, is additionally a significant contributor to antibiotic resistance (9). Given the role of recombination in antibiotic resistance evolution, increasing recombination rates by administering drugs like those studied here will almost certainly accelerate resistance emergence. Altogether, our study helps understand the underpinnings of antibiotic resistance evolution and entails important considerations for diverse fields, from evolution to biomedicine or cancer biology.

## Material and methods

### Bacteria and growth conditions

The primary bacterial strain in this study was the *Escherichia coli* MG1655 transformed with the plasmid pRHOMO recombination system. We also used a *recA*-deficient derivative of the strain MG1655 (17). To determine the impact of antimicrobials on the generation of DSBs, the *E. coli* strain SMR14354 was used (25). The *E. coli* strain DH-5 alpha [*fhuA2 lac(del)U169 phoA glnV44 Φ80’ lacZ(del)M15 gyrA96 recA1 relA1 endA1 thi-1 hsdR17*] was used to maintain the plasmid of recombination system pRHOMO (17). All strains were routinely cultured in Lysogeny Broth (LB medium, Carl Roth, Germany), supplemented with antibiotics when appropriate. For solid cultures, we added bacteriological agar to LB at a final concentration of 1.6%.

### Minimal Inhibitory Concentration

The minimal inhibitory concentration (MIC) was performed by broth microdilution method according to EUCAST guidelines, except that LB medium was used instead of Mueller-Hinton broth. The antibiotics used were ciprofloxacin, mitomycin C, bleomycin, trimethoprim, streptomycin, kanamycin, chloramphenicol, ceftazidime, cefotaxime, fosfomycin and rifampicin (Supplementary Table S1).

### Recombination frequency

A genetic assay was carried out as previously described to estimate the impact of antimicrobials on recombination frequency (17). The system consisted of a plasmid harboring two truncated *tetA* alleles separated by an antibiotic resistance cassette (*aacC1*, conferring gentamicin resistance, Supplementary Figure S1). Recombination restores the functional *tetA* gene, thereby conferring tetracycline resistance, which can be selected for (13, 17). Thus, this assay allows quantification of the recombination frequency with high sensitivity due to the elevated copy number of the plasmid. We exposed 1 mL of culture containing approximately 1-2×10^7^ bacteria [dilution of 1/10 of mid-exponential culture containing 1×10^8^ colony-forming units per millilitre (CFU/mL)] to different concentrations of different antimicrobials (to match for all of them their respective MIC values) for three hours, similarly to a previously described experiment [6]. After incubation with gentle shaking, we added 9 mL of fresh LB to the cultures and centrifuged them at 4000 x g for 10 minutes to reduce and remove the antimicrobials. The pellets were washed twice with 2 mL of LB, and cells were allowed to recover for one hour before adding ampicillin (to maintain the plasmid) to a final concentration of 50 µg/mL and incubated overnight with shaking at 37°C. The next day, appropriate dilutions were plated in LB agar containing 50 µg/mL of ampicillin to estimate the viability and LB agar containing 30 µg/mL of tetracycline to estimate the number of recombinants. Each experiment consisted of five replicas and was repeated twice. Recombinant frequencies were expressed as the ratio of means of recombinant number by the number of viable bacteria.

### Influence of antimicrobials on the generation of double-strand breaks

The strain SMR14354 (25) was used to measure the number of double-strand breaks when the strain is incubated with different concentrations of tested antimicrobials. This strain carries a reporter fusion of the protein Gam, able to detect double DNA ends and a green fluorescent protein (GFP). The strain also harbours a unique I-*SceI* restriction site in the chromosome and carries the gene for the restriction enzyme *SceI* whose expression is regulated by an arabinose-inducible promoter [20]. Mid exponential cultures containing 0.5×10^8^ CFU/mL were exposed to different concentrations of selected antibiotics for four hours. Then, 5 µl drops of the cultures were added to LB agar blocks containing 1 µM Propidium Iodide (PI, Sigma, Germany) and the covered bacterial area of the blocks were placed on a circular bottom glass open cultivation system II (PeCon GmbH, Germany). This device consists of a metallic holder with a 0.17 mm round-bottom glass and a glass lid that limits evaporation. The device was placed on a robotic inverted microscope Nikon Eclipse T2 (Nikon, Japan). Several fields of each sample were taken automatically using the red and green channels to capture the signal of PI (as viability criteria) and the signal of Gam-GFP fusion to detect DSBs, respectively. Three independent overnight cultures were analyzed per antimicrobial per concentration.

### Influence of iron on the level of double-strand breaks and recombination frequency of bleomycin-treated cells

Double-strand breaks induced by bleomycin depend on intracellular iron concentration (20). To understand the relationship between double-strand breaks and recombination frequency, bleomycin-treated cells were subjected to iron depletion. The *E. coli* MG1655 carrying the pRHOMO construct cultures treated with 1X MIC of bleomycin alone and bleomycin plus 2-2′ bipyridyl (50 µM) were examined for changes in recombination rate and the number of double-strand breaks determined as described elsewhere in this section.

### Influence of initial population sizes on recombination frequency

To determine the impact of different initial population sizes on the recombination frequency, different sets of cultures of *E. coli* MG1655-pRHOMO, with inoculum sizes ranging from 10^4^ to 10^8^ CFU/mL, were inoculated in 10 mL of LB supplemented with ampicillin to a final concentration of 50 µg/mL and incubated overnight with shaking at 37°C. The next day, dilutions were plated in LB agar containing 50 µg/mL of ampicillin to estimate the viability and in LB agar containing 30 µg/mL of tetracycline to estimate the number of recombinants. Each experiment consisted of five replicas. Recombinant frequencies were expressed as the ratio of means of recombinant number by the number of viable bacteria.

### Experimental evolution of ciprofloxacin-resistant *Escherichia coli* MG1655

The *E. coli* MG1655 strain used in all experiments served as the starting population for this experimental evolution study. Preadaptation was performed in triplicate by inoculating 50 ml Falcon tubes with 3.7 ml of LB medium containing single colonies (preserving similar air/volume ratio interface of 96 multiwell plates). The cultures were incubated for 24 h at 37 °C without shaking and then transferred daily for three days to fresh medium (dilution 1:1000). The optical density (OD) at 600 nm was measured using an NP80 NanoPhotometer (Implen GmbH, Munich, Germany) and compared to that of the original strain. The replicate line with the OD most similar to the original strain was cryopreserved and used as a preadapted strain for subsequent experiments.

For the selection experiment, five replicates were prepared from individual colonies of the pre-adapted ancestor, along with one unselected control line. The cultures were grown to the mid-exponential growth phase, and 10 µL of each was inoculated into 96-well microtiter plates (Greiner Bio-One International GmbH, Kremsmünster, Austria) containing 200 µL of LB medium (VWR International, Darmstadt, Germany) with ciprofloxacin (Merck KGaA, Darmstadt, Germany) at half the minimum inhibitory concentration (MIC = 0.03125 µg/mL). The plates were incubated at 37 °C and shaken every 30 min for 5 s in a BioTek Synergy HTX plate reader (BioTek, Santa Clara, USA).

The cultures were passated daily in fresh medium at a 1:100 dilution. The ciprofloxacin concentration was maintained or doubled once the optical density (OD) of the treated cultures reached at least half that of the positive control. The experiment was terminated after six passes at four times the minimum inhibitory concentration (MIC). Three of the five replicates survived; these were cryopreserved and subjected to MIC determination. Both the resulting populations (p) and the isolated single colonies (c) were stored for phenotypic and genomic analyses. All isolates were preserved at −80 °C in LB medium with 20% glycerol.

Whole-genome sequencing of populations (p) and isolates (c) was performed by MicrobesNG (Birmingham, UK) using the Illumina NovaSeq 6000 platform (Illumina, San Diego, USA) with a 250 bp paired-end protocol at 30-fold coverage, according to MicrobesNG’s submission procedures. Three clones with different mutations and different ciprofloxacin resistance were selected for recombination experiments. The strains were transformed by electroporation with the plasmid pRHOMO and the recombination frequencies were determined as previously described.

### Statistical analysis

To compare experimental groups with their control, the Welch test was performed. To avoid biases introduced by multiple corrections due to comparison against control, each antimicrobial experiment had its own non-treated control. In addition, the analysis of covariance (ANCOVA) was carried out to estimate the co-occurrence of double-strand breaks and increase in recombination rate induced by different antibiotics. P-values less than or equal to 0.05 were considered statistically significant. Statistical tests and graph plotting were performed with R (42) and GraphPad Prism version 9.00 (GraphPad Software, Inc., San Diego, CA).

## Data Availability

All relevant data are provided the manuscript and its supporting Information files, including raw data.

## Acknowledgements

We are especially grateful to Sophie Armitage, Audrey Menegaz Proença (Freie Universität Berlin), Luís M Silva (University of Neuchâtel) and Céline Vidaillac (Oxford University Clinical Research Unit, OUCRU) for critical comments and proofreading. We thank Jerónimo Rodríguez-Beltrán from the Spanish National Centre for Biotechnology (Spanish National Research Council, CSIC) for providing us with the construct to determine recombination frequency in *E. coli* and Susan M. Rosenberg from Baylor College of Medicine (Texas, USA) for the strain of *E. coli* SMR14354 to visually determine double-strand breaks. We are grateful to Rees Kassen (University of Ottawa) for a scientific discussion on recombination in bacteria and antibiotics, which allowed us to develop the central hypothesis of the current study.

## Author contributions

A.R.R. conceived and designed the concept of the experiments. A.N., D.R. and N.P. performed the experiments and conducted data analysis. A.N., D.R., J.R., J.B. and A.R.R co-wrote and proofread the manuscript. J.R., J.B. and A.R.R. contributed with resources and funding. All the authors discussed, commented and agreed on the manuscript.

## Supplementary material

**Figure S1.**
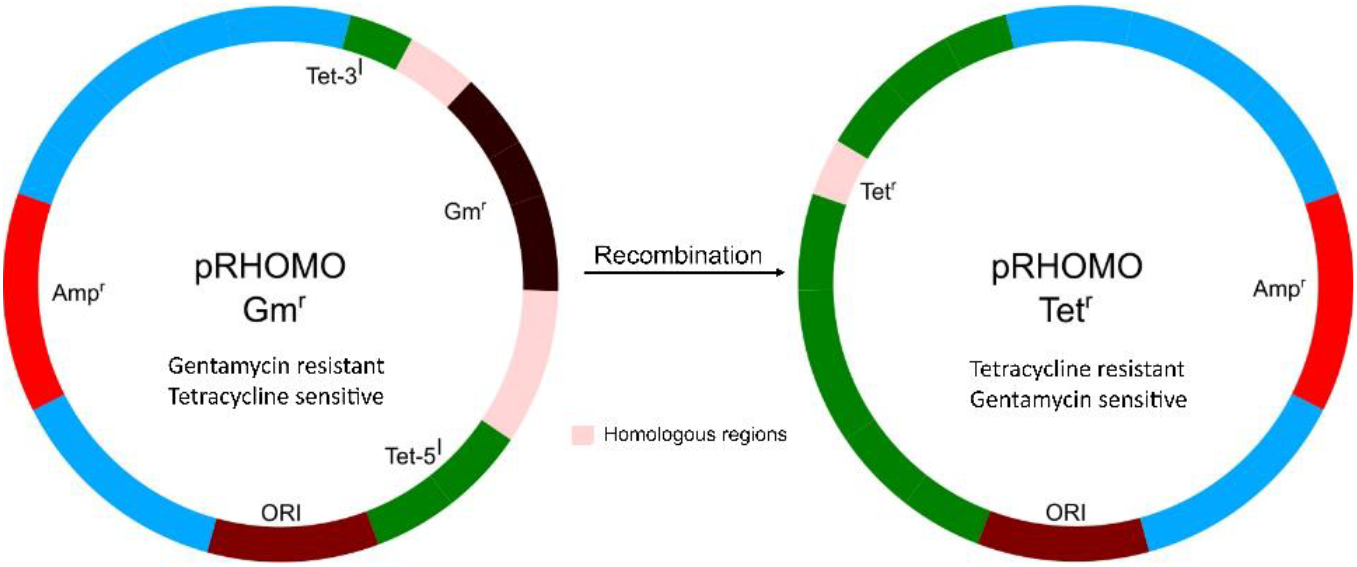
Representation of the plasmid system to measure the recombination frequency (modified from reference (16). **The system consists of a plasmid harbouring two truncated** *tetA* alleles separated by an antibiotic resistance cassette (*aacC1*, conferring resistance to gentamycin). Recombination restores the functional *tetA* gene, thereby restoring a functional tetracycline-resistant cassette, which can be selected on an agar plate containing tetracycline.

**Figure S2.**
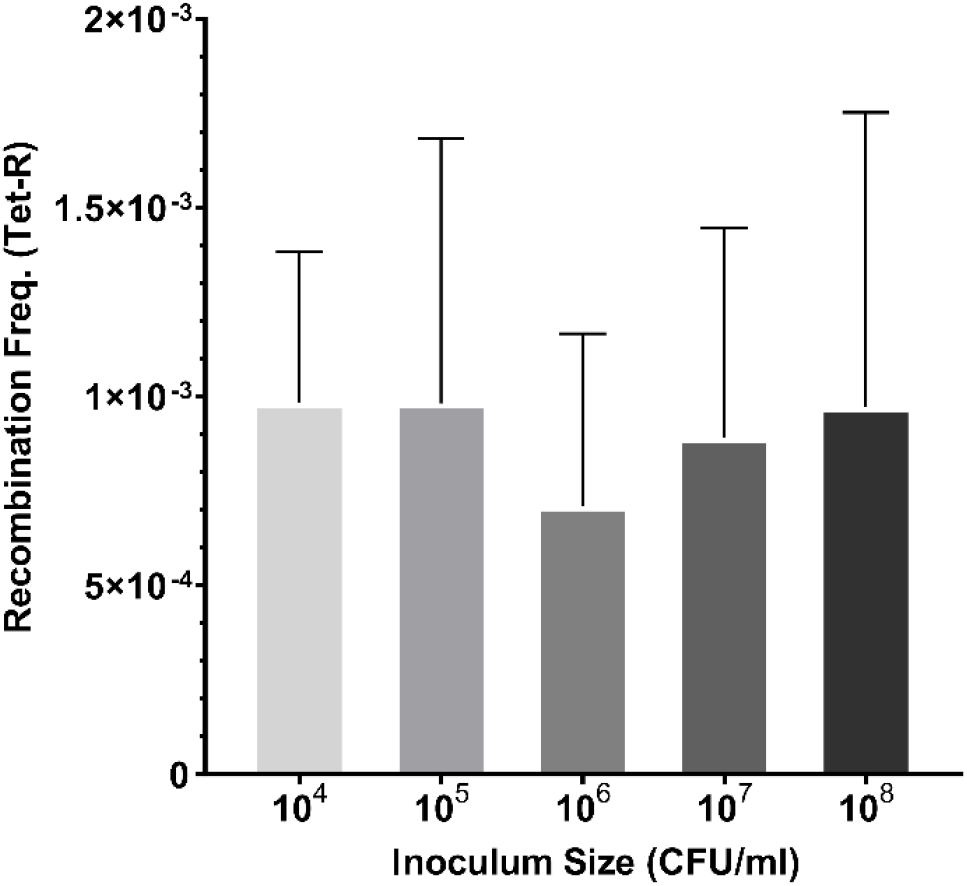
Effect of population size on recombination frequency. Cultures *E. coli* MG1655 harbouring the plasmid pRHOMO were inoculated with growing bacterial numbers ranging from 10^4^ to 10^8^ bacteria (Figure S2). There is no statistical difference in recombination frequency due to the additional number of divisions (Analysed by one-way ANOVA followed by Bonferroni’s multiple comparisons tests with a single pooled variance). Each experiment consisted of five replicates. The bars represent the means, while the error bars represent the standard deviations (n=5 + SD).

**Table S1.**
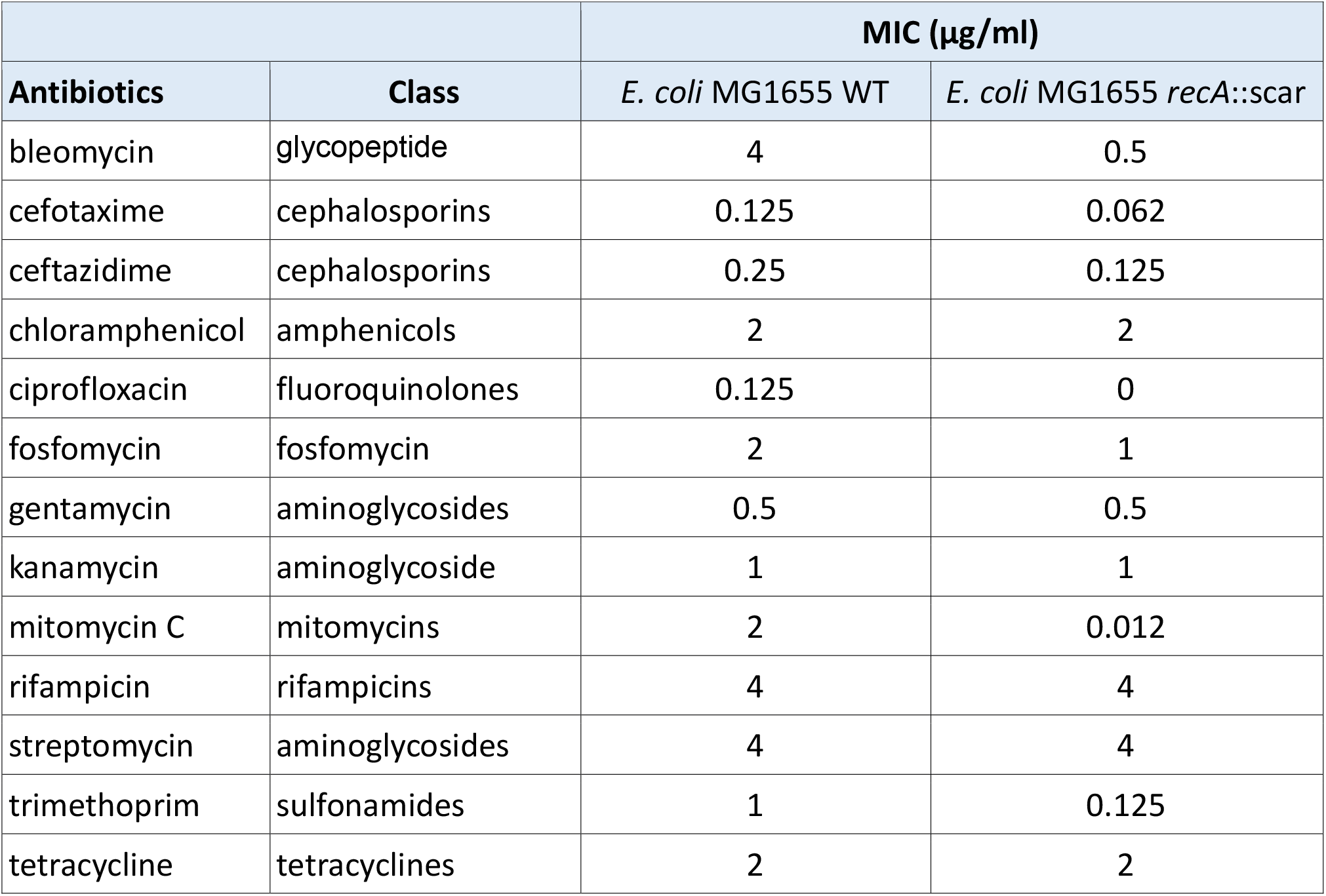
Minimal inhibitory concentration (MIC) values for *E. coli* MG1655 and its derivative mutant *E. coli* MG1655 *recA*::scar for different antimicrobials used in this work.

| Antibiotics | Class | MIC (µg/ml) |  |
| --- | --- | --- | --- |
|  |  | *E. coli* MG1655 WT | *E. coli* MG1655 *recA::scar* |
| bleomycin | glycopeptide | 4 | 0.5 |
| cefotaxime | cephalosporins | 0.125 | 0.062 |
| ceftazidime | cephalosporins | 0.25 | 0.125 |
| chloramphenicol | amphenicols | 2 | 2 |
| ciprofloxacin | fluoroquinolones | 0.125 | 0 |
| fosfomycin | fosfomycin | 2 | 1 |
| gentamycin | aminoglycosides | 0.5 | 0.5 |
| kanamycin | aminoglycoside | 1 | 1 |
| mitomycin C | mitomycins | 2 | 0.012 |
| rifampicin | rifampicins | 4 | 4 |
| streptomycin | aminoglycosides | 4 | 4 |
| trimethoprim | sulfonamides | 1 | 0.125 |
| tetracycline | tetracyclines | 2 | 2 |

## References

1. Naghavi et al. 2024. Global burden of bacterial antimicrobial resistance 1990–2021: a systematic analysis with forecasts to 2050. The Lancet 404:1199–1226.

2. Baker S. 2015. A return to the pre-antimicrobial era? Science 347:1064–1066.

3. Murray CJ, et al. 2022. Global burden of bacterial antimicrobial resistance in 2019: a systematic analysis. The Lancet 0.

4. Rodríguez-Rojas A, Rodríguez-Beltrán J, Couce A, Blázquez J. 2013. Antibiotics and antibiotic resistance: A bitter fight against evolution. International Journal of Medical Microbiology 303:293–297.

5. Zhang D, Cui Y, Zhang X. 2019. Estimating factors related to fluoroquinolone resistance based on one health perspective: Static and dynamic panel data analyses from Europe. Frontiers in Pharmacology 10:1145.

6. Witzany C, Bonhoeffer S, Rolff J. 2020. Is antimicrobial resistance evolution accelerating? PLoS pathogens 16.

7. Hooper DC. 1998. Bacterial topoisomerases, anti-topoisomerases, and anti-topoisomerase resistance. Clinical infectious diseases: an official publication of the Infectious Diseases Society of America 27 Suppl 1.

8. Hooper DC, Jacoby GA. 2016. Topoisomerase Inhibitors: Fluoroquinolone Mechanisms of Action and Resistance. Cold Spring Harbor Perspectives in Medicine 6.

9. Blázquez J, Couce A, Rodríguez-Beltrán J, Rodríguez-Rojas A, Blazquez J, Couce A, Rodriguez-Beltran J, Rodriguez-Rojas A. 2012. Antimicrobials as promoters of genetic variation. Curr Opin Microbiol2012/08/15. 15:561–569.

10. Gutierrez A, Laureti L, Crussard S, Abida H, Rodríguez-Rojas A, Blázquez J, Baharoglu Z, Mazel D, Darfeuille F, Vogel J, Matic I. 2013. β-Lactam antibiotics promote bacterial mutagenesis via an RpoS-mediated reduction in replication fidelity. Nature communications 4:1610.

11. Nordman J, Wright A. 2008. The relationship between dNTP pool levels and mutagenesis in an Escherichia coli NDP kinase mutant. Proc Natl Acad Sci U S A2008/07/16. 105:10197– 10202.

12. Rodríguez-Rojas A, Makarova O, Rolff J. 2014. Antimicrobials, stress and mutagenesis. PLoS pathogens 10:e1004445.

13. Rodríguez-Rojas A, Moreno-Morales J, Mason AJ, Rolff J. 2018. Cationic antimicrobial peptides do not change recombination frequency in Escherichia coli. Biology Letters 14:20180006.

14. Sandegren L, Andersson DI. 2009. Bacterial gene amplification: implications for the evolution of antibiotic resistance. Nature Reviews Microbiology 7:578–588.

15. Elliott KT, Cuff LE, Neidle EL. 2013. Copy number change: evolving views on gene amplification. Future Microbiology 8:887–899.

16. Lopez E, Elez M, Matic I, Blazquez J, López E, Elez M, Matic I, Blázquez J. 2007. Antibiotic-mediated recombination: ciprofloxacin stimulates SOS-independent recombination of divergent sequences in Escherichia coli. Molecular microbiology2007/03/23. 64:83–93.

17. Rodríguez-Beltrán J, Tourret J, Tenaillon O, López E, Bourdelier E, Costas C, Matic I, Denamur E, Blázquez J. 2015. High Recombinant Frequency in Extraintestinal Pathogenic Escherichia coli Strains. Molecular biology and evolution 32:1708–16.

18. López E, Blázquez J, Lopez E, Blazquez J. 2009. Effect of subinhibitory concentrations of antibiotics on intrachromosomal homologous recombination in Escherichia coli. Antimicrob Agents Chemother2009/06/03. 53:3411–3415.

19. Kenyon CJ, Walker GC. 1980. DNA-damaging agents stimulate gene expression at specific loci in Escherichia coli. Proc Natl Acad Sci U S A1980/05/01. 77:2819–2823.

20. Xu T, Brown W, Marinus MG. 2012. Bleomycin Sensitivity in Escherichia coli is Medium-Dependent. PLoS ONE 7:e33256.

21. Do Thi T, López E, Rodríguez-Rojas A, Rodríguez-Beltrán J, Couce A, Guelfo JRR, Castañeda-García A, Blázquez J, Thi TD, Lopez E, Rodriguez-Rojas A, Rodriguez-Beltran J, Couce A, Guelfo JRR, Castaneda-Garcia A, Blazquez J. 2011. Effect of recA inactivation on mutagenesis of Escherichia coli exposed to sublethal concentrations of antimicrobials. J Antimicrob Chemother2011/01/08. 66:531–538.

22. Blázquez J, Rodríguez-Beltrán J, Matic I. 2018. Antibiotic-Induced Genetic Variation: How It Arises and How It Can Be Prevented. Annual Review of Microbiology. Annual Reviews Inc. 10.1146/annurev-micro-090817-062139.

23. Keller KL, Overbeck-Carrick TL, Beck DJ. 2001. Survival and induction of SOS in Escherichia coli treated with cisplatin, UV-irradiation, or mitomycin C are dependent on the function of the RecBC and RecFOR pathways of homologous recombination. Mutation research 486:21–29.

24. Taddei F, Vulic M, Radman M, Matic I. 1997. Genetic variability and adaptation to stress. EXS1997/01/01. 83:271–290.

25. Shee C, Cox BD, Gu F, Luengas EM, Joshi MC, Chiu L-Y, Magnan D, Halliday JA, Frisch RL, Gibson JL, Nehring RB, Do HG, Hernandez M, Li L, Herman C, Hastings PJ, Bates D, Harris RS, Miller KM, Rosenberg SM. 2013. Engineered proteins detect spontaneous DNA breakage in human and bacterial cells. eLife 2:e01222.

26. Chen J, Ghorai MK, Kenney G, Stubbe J. 2008. Mechanistic studies on bleomycin-mediated DNA damage: multiple binding modes can result in double-stranded DNA cleavage. Nucleic acids research 36:3781–90.

27. Rodríguez-Rojas A, Makarova O, Müller U, Rolff J. 2015. Cationic Peptides Facilitate Iron-induced Mutagenesis in Bacteria 11:e1005546.

28. Povirk LF, Wübter W, Köhnlein W, Hutchinson F. 1977. DNA double-strand breaks and alkali-labile bonds produced by bleomycin. Nucleic acids research 4:3573–80.

29. Jang S, Imlay JA. 2007. Micromolar intracellular hydrogen peroxide disrupts metabolism by damaging iron-sulfur enzymes. J Biol Chem2006/11/15. 282:929–937.

30. Datsenko KA, Wanner BL. 2000. One-step inactivation of chromosomal genes in Escherichia coli K-12 using PCR products. Proc Natl Acad Sci U S A2000/06/01. 97:6640–6645.

31. Chaveroche MK, Ghigo JM, d’Enfert C. 2000. A rapid method for efficient gene replacement in the filamentous fungus Aspergillus nidulans. Nucleic Acids Research 28:e97.

32. Kono M, Saitoh Y, Kasai M, Shirahata K. 1995. Isolation of albomitomycins from solutions of 7-amino substituted mitomycins; mitomycin C and KW-2149. The Journal of antibiotics 48:179–81.

33. Umezawa H, Maeda K, Takeuchi T, Okami Y. 1966. New antibiotics, bleomycin A and B. The Journal of antibiotics 19:200–9.

34. May JM, Owens TW, Mandler MD, Simpson BW, Lazarus MB, Sherman DJ, Davis RM, Okuda S, Massefski W, Ruiz N, Kahne D. 2017. The antibiotic novobiocin binds and activates the ATPase that powers lipopolysaccharide transport. Journal of the American Chemical Society 139:17221.

35. Riaz L, Mahmood T, Khalid A, Rashid A, Ahmed Siddique MB, Kamal A, Coyne MS. 2018. Fluoroquinolones (FQs) in the environment: A review on their abundance, sorption and toxicity in soil. Chemosphere 191:704–720.

36. Millanao AR, Mora AY, Villagra NA, Bucarey SA, Hidalgo AA. 2021. Biological Effects of Quinolones: A Family of Broad-Spectrum Antimicrobial Agents. Molecules (Basel, Switzerland) 26:7153.

37. Bolzán AD, Bianchi MS. 2018. DNA and chromosome damage induced by bleomycin in mammalian cells: An update. Mutation Research/Reviews in Mutation Research 775:51–62.

38. Laehnemann D, Peña-Miller R, Rosenstiel P, Beardmore R, Jansen G, Schulenburg H. 2014. Genomics of Rapid Adaptation to Antibiotics: Convergent Evolution and Scalable Sequence Amplification. Genome Biology and Evolution 6:1287.

39. Chu H-Y, Sprouffske K, Wagner A. 2017. The role of recombination in evolutionary adaptation of Escherichia coli to a novel nutrient. Journal of Evolutionary Biology 30:1692– 1711.

40. D’Aeth JC, van der Linden MPG, McGee L, de Lencastre H, Turner P, Song JH, Lo SW, Gladstone RA, Sá-Leão R, Ko KS, Hanage WP, Breiman RF, Beall B, Bentley SD, Croucher NJ. 2021. The role of interspecies recombination in the evolution of antibiotic-resistant pneumococci. eLife 10.

41. Lawrence JG, Retchless AC. 2009. The Interplay of Homologous Recombination and Horizontal Gene Transfer in Bacterial Speciation, p. 29–53. In Methods in molecular biology (Clifton, N.J.).

42. R Core Team. 2017. R: A language and environment for statistical computing. R Foundation for Statistical Computing, Vienna, Austria.

